# Proposed Mechanism for Monomethylarsonate Reductase Activity of Human Omega-class Glutathione Transferase GSTO1-1

**DOI:** 10.1101/2021.11.07.467205

**Authors:** Aaron J. Oakley

## Abstract

Contamination of drinking water with toxic inorganic arsenic is a major public health issue. The mechanisms of enzymes and transporters in arsenic elimination are therefore of interest. The human omega-class glutathione transferases have been previously shown to possess monomethylarsonate (V) reductase activity. To further understanding of this activity, molecular dynamics of human GSTO1-1 bound to glutathione with a monomethylarsonate isostere were simulated to reveal putative monomethylarsonate binding sites on the enzyme. The major binding site is in the active site, adjacent to the glutathione binding site. Based on this and previously reported biochemical data, a reaction mechanism for this enzyme is proposed. Further insights were gained from comparison of the human omega-class GSTs to homologs from a range of animals.

## Introduction

Chronic exposure to arsenic is associated with several conditions including skin lesions; peripheral neuropathy; and cancer (Ratnaike, 2003). Contamination of drinking-water with soluble inorganic arsenic is a significant problem globally. It is estimated that that the drinking water of at least 140 million people in 50 countries contains arsenic at levels above the WHO provisional guideline (10 μg/L) (Ravenscroft et al., 2009). The relationship between chronic consumption of arsenic and disease is poorly understood.

Investigation of arsenic metabolism in humans has revealed a series of enzymes and transporters that reduce and export arsenicals from cells. However, the pathway(s) of arsenic metabolism remain controversial. In one scheme (“oxidative methylation”), arsenite (As^III^(OH)_3_) is converted to monomethylarsonic acid (CH_3_As^V^O(OH)_2_; MMA^V^) and dimethylarsinic acid ((CH_3_)_2_As^V^O(OH); DMA^V^) (Aposhian, 1997). An alternative pathway (“reductive methylation”) was proposed that involves the non-enzymatic production of arsenic triglutathione from arsenite and subsequent methylation by arsenic methyltransferase (Hayakawa et al., 2005). Arsenic metabolism and toxicity has been reviewed recently (Khairul et al., 2017).

In 2001, the human MMA^V^ reductase (E.C. 1.20.4.2) was isolated and shown to be identical to glutathione transferase omega-1 (GSTO1-1) (Zakharyan et al., 2001), an enzyme belonging to the cytosolic glutathione transferase (GST) superfamily of enzymes that use glutathione (GSH, γ-glutamyl-cysteinyl-glycine) in a diverse range of detoxification reactions (Oakley, 2011). Reduction of MMA^V^ is the rate-limiting step in the biotransformation of inorganic arsenic (Zakharyan et al., 2001). The MMA^V^ reducing activity of liver cytosol from GSTO1 KO mice has been shown to be 20% of that of wild-type mice (Chowdhury et al., 2006).

In humans, two genes (*GSTO1* and *GSTO2*) encode two omega-class GST isozymes that share 64% sequence identity (Whitbread et al., 2005). Both human GSTO1-1 and GSTO2-2 isozymes catalyse the reduction of pentavalent methylated arsenic species (Schmuck et al., 2005, Zakharyan et al., 2001). Several polymorphisms in the human omega-class genes have been identified. The GSTO1-A140D polymorphism is the most extensively studied, having been linked to cancer in arsenic-exposed populations (Beebe-Dimmer et al., 2012). The GSTO1-A140D and E208K polymorphisms have been implicated in increased risk of apoptosis and inflammatory-related diseases in arsenic-exposed populations (Escobar-García et al., 2012).

The first crystal structure of human GSTO1 (hGSTO1-1) at 2.0 Å resolution revealed a homodimeric enzyme whose fold is typical of the cytosolic GSTs (Board et al., 2000). The structure of human GSTO2 (hGSTO2-2) has broadly similar features to hGSTO1-1 (Zhou et al., 2012). Mammalian cytosolic GSTs have been divided into 7 evolutionarily distinct classes: alpha, mu, pi, sigma, theta, zeta, and omega (Board et al., 2000). GSTs (EC 2.5.1.18) catalyse the conjugation of xenobiotics with electrophilic centres to GSH and other reactions including degradation of tyrosine and biosynthetic reactions generating leukotrienes, prostaglandins, progesterone and testosterone (Hayes et al., 2005). All mammalian cytosolic GSTs are dimeric. Each monomer has an N-terminal domain adopting the thioredoxin-fold that binds GSH (the “G-site”) and a C-terminal domain composed of α-helices that binds a co-substrate (called the “H-site”, after the hydrophobic nature of the co-substrate in several GST classes). Two major subgroups have been identified within the cytosolic GSTs based on sequence and structural similarity (Atkinson and Babbitt, 2009): the Y-GSTs, which utilize a tyrosine residue to activate GSH, and the S/C-GSTs that utilize serine or cysteine residue. Several lines of evidence suggest that in GSTs containing a catalytic tyrosine or serine residue, the OH group donates a hydrogen bond to GSH, lowering the p*K*a and stabilizing the thiolate anion (Oakley, 2011). In the set of known human cytosolic GSTs, only the omega-class enzymes utilize a catalytic cysteine residue (C32 in hGSTO1-1 and hGSTO2-2) (Whitbread et al., 2003). In this regard, the omega-class GSTs resemble the glutaredoxins, and like the glutaredoxins they can catalyse GSH-dependent reductase reactions. In addition to MMA^V^, substrates reduced by the omega-class GSTs include dehydroascorbate (Whitbread et al., 2003). Like the glutaredoxins, hGSTO1-1 and hGSTO2-2 can form mixed-disulfides with GSH (Zhou et al., 2012).

Surprisingly, the bacterial lipopolysaccharide (LPS)-stimulated inflammatory responses were shown to be regulated by hGSTO1 (Menon et al., 2014). Specifically, hGSTO1 regulates NLRP3 inflammasome activation *via* deglutathionylation of serine/threonine-protein kinase NEK7 (Hughes et al., 2019). As such, human GSTO1 is a target for inhibition for the treatment of a variety of inflammatory disease states. Several GSTO1 inhibitors have been reported that covalently modify C32 (Tsuboi et al., 2011, Dai et al., 2019, Xie et al., 2020).

Attempts to observe MMA^V^ binding in GSTO1-1 by X-ray crystallography have been unsuccessful to date. To gain insights into the MMA^V^ reductase activity of GSTO1-1, sub-microsecond timescale molecular dynamics trajectories were calculated with the enzyme and monomethylphosphonate as an MMA^V^ isostere. Methylphosphonate and methylarsonate both have tetrahedral geometry and similar p*K*_a_ values (p*K*_a1_ = 2.12; p*K*_a2_ = 7.29 and p*K*_a1_ = 3.6; p*K*_a2_ = 8.2 respectively). To enhance sampling of ligand binding, a low-mass MD (LMD) approach was taken whereby the masses of the heavy atoms of MMP were set to 10% of their normal atomic weights, thereby accelerating diffusion and binding/unbinding events. Similar approaches have been used successfully to study protein folding (Pang, 2014).

## Methods

MD trajectories were calculated by NAMD 2.14 (Phillips et al., 2020) using the OPLS-AA/M all-atom force field (Robertson et al., 2015). Based on the MMA^V^ p*K*_a_ values, the monovalent anion was selected as the most relevant species at physiological pH. Parameters for the isostere monomethylphosphate (MMP^V^) were generation by the LigParGen server (Dodda et al., 2017). The simulated systems consisted of a GSTO1-1 dimer, 38 MMP ions (~100 mM) and water molecules in a 90 × 90 × 90 Å cube. Sufficient sodium ions were added to give the system zero net charge. All simulations were run as NpT ensembles (temperature, 310 K; pressure, 101.325 kPa) with periodic boundary conditions. Temperature was controlled using Langevin dynamics (damping constant 5 ps^−1^). Pressure control used the Nosé-Hoover Langevin piston (period 100 fs; decay rate 50 fs). A multiple time-step approach was used with 1, 2, and 4 fs for bonded, non-bonded and long-range electrostatic calculations respectively. The Particle-mesh Ewald with a grid resolution of ~1 Å was used to calculate long-range electrostatic forces. van der Waals’ interactions were smoothly scaled to zero between 10 and 12 Å. To prevent excessive rotation/translation of the enzyme, weak harmonic restraints (0.01 kcal mol^−1^ Å^−2^) were applied to Cα atoms. The system was subjected to energy minimization (10,000 steps) prior to equilibration. Coordinates were saved every 10 ps for analysis. The total simulation time was 0.5 μs (500 × 10^6^ steps).

Trajectory data were analysed in VMD (Humphrey et al., 1996). Volumetric density maps with grid spacing of 0.5 Å were calculated by replacing all phosphorus atoms with a normalized gaussian distribution with standard deviation equal to the atomic radius at each grid point. The final map represents the average density over all frames of the trajectory. Since GSTO1-1 is a homodimer, sampling was further increased by averaging the density around each monomer (analogous to NCS averaging in crystallography).

The active site of human GSTO1 was compared with GSTO2 and homologs of other species. Models of GSTO1 homologs predicted by AlphaFold (Jumper et al., 2021) were obtained from the European Bioinformatic Institute database corresponding to the following species (and UniProt IDs): Mouse (O09131), Rat GSTO1 (Q9Z339) *Drosophila melanogaster* (Q9VSL6) and *C. elegans* (P34345). Sequences of selected GSTO1 homologs were used to generate a sequence alignment in ClustalW (Larkin et al., 2007). The Adaptive Poisson-Boltzmann Solver (APBS) software (Jurrus et al., 2018) was used to compute electrostatic potential maps.

## Results

The simulated system contained 67,589 atoms, including 7,736 protein, 72 GSH, 342 MMP and 59,397 water. The MD trajectory contained 50,000 frames. Atom density maps are shown in Figure 1. The maps prior to averaging densities in both monomers (Figure 1A) shows that the region favourable to ligand binding are consistent between the two monomers. The density map after averaging is shown for the monomer in Supplementary Figure 1. The region with the highest density peaks (the active site) is shown in Figure 1B. Two favourable binding loci are observed. Sample trajectory frames with MMP bound at these loci show a close interaction of the ligand with the guanidinium group of R132, placing the ligand 9 Å and 5 Å from the GSH sulhydryl group, corresponding to density maxima of 0.31 and 0.25 Da Å^−3^ respectively (Figure 1C, D). For reference, the average density in the solvent region is approximately 0.0015 Da Å^−3^.

**Figure 1.**
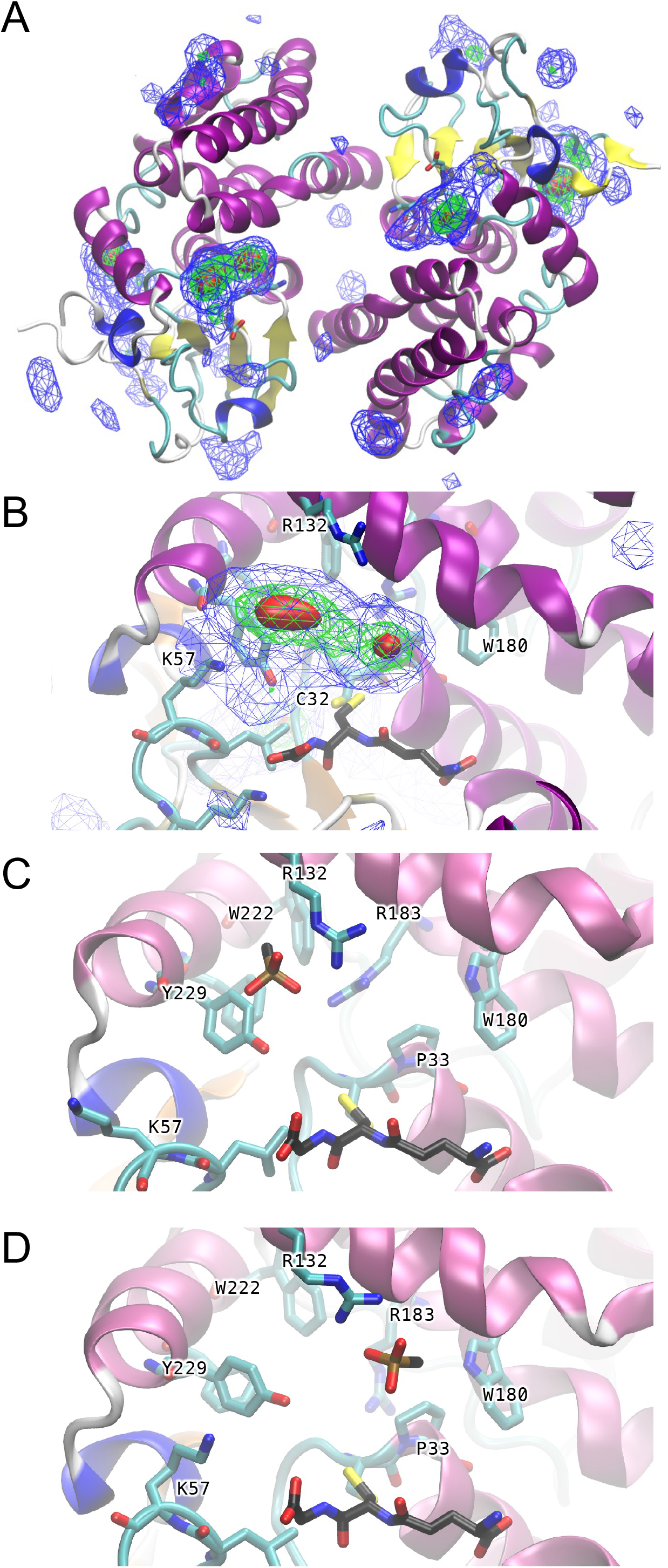
Density isosurfaces for P atoms in blue, green and red are contoured at 0.02, 0.1 and 0.2 Da Å^−3^ respectively. (A) Cartoon representation of GSTO1-1 showing density isosurfaces prior to symmetry-based averaging. (B) Active site of GSTO1 with density isosurfaces averaged over both monomers. GSH is shown in stick form with black carbon atoms. (C) and (D) snapshots from the MD trajectory showing MMP at the two density maxima shown in (B).

A third (“accessory”) locus for MMP binding was observed to the side of the β-sheet near the end of strand β2 (Supplementary Figure 1C). This is near binding sites for sulfate in several GSTO1-1 crystal structures (e.g. PDB ID 5YVN; (Saisawang et al., 2019). The density maximum here is 0.23 Da Å^−3^. The side is bound by residues R30, R39, K43 and H49 (Supplementary Figure 1C).

During the simulation, rotations about the χ_1_ angle of the GSH cysteine were observed, resulting in the movement of the Sγ atom away from the C32-SγH group and orienting it toward the Y229 Oη as shown in Figures 1C and 1D. During the simulation, the GSH sulfydryl to Y229 Oη distance (mean ± standard deviation) was 5.6 ± 1.1 Å, with a minimum observed distance of 2.9 Å.

The H-sites of human GSTO1 and GSTO2 are shown in Figure 2, together with models of homologs from mouse, rat, *Drosophila* and *Caenorhabditis*. Except for some surface loops and termini regions, the bulk of the models (including active site residues) had pLDDT values > 90 (indicating “very high confidence”) and were judged suitable for structure comparisons. The sequence alignment these GSTO homologs plus those from cow and pig is shown in Figure 3.

**Figure 2.**
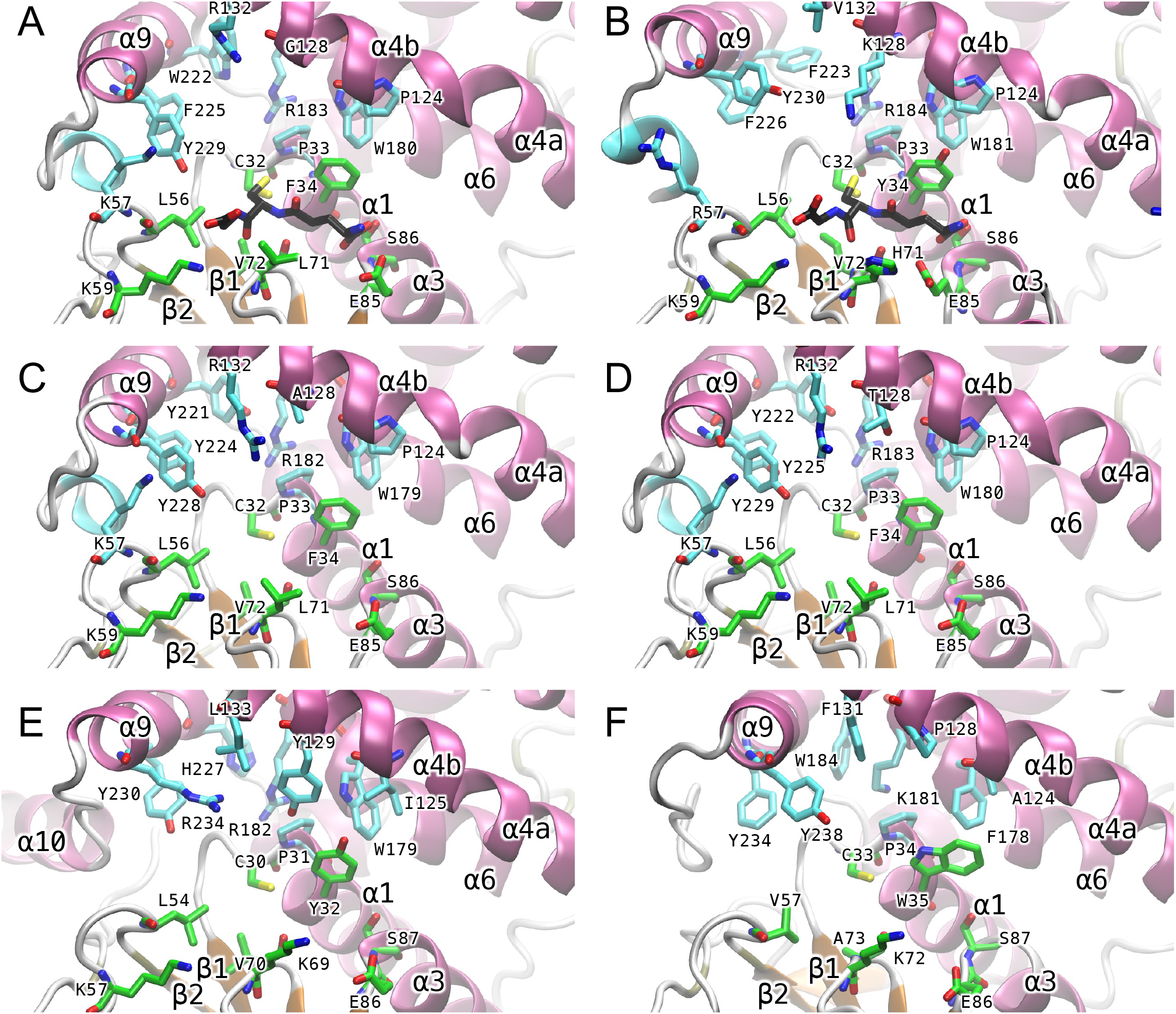
Active sites of GSTO homologs. Crystal structures of (A) human GSTO1 (PDB ID 1EEM) (B) human GSTO2 (PDB ID 3Q19) and AlphaFold models of (C) mouse GSTO1, (D) rat GSTO1, (E) *D. melanogaster* GSTO1, and (F) *C. elegans* GSTO1. A cartoon representation is used for the backbone. GSH in human GSTO crystal structures is in stick form (carbon atoms black). Sidechains for G-site (carbon atoms green) and H-site (carbon atoms cyan) residues are also in stick form.

**Figure 3.**
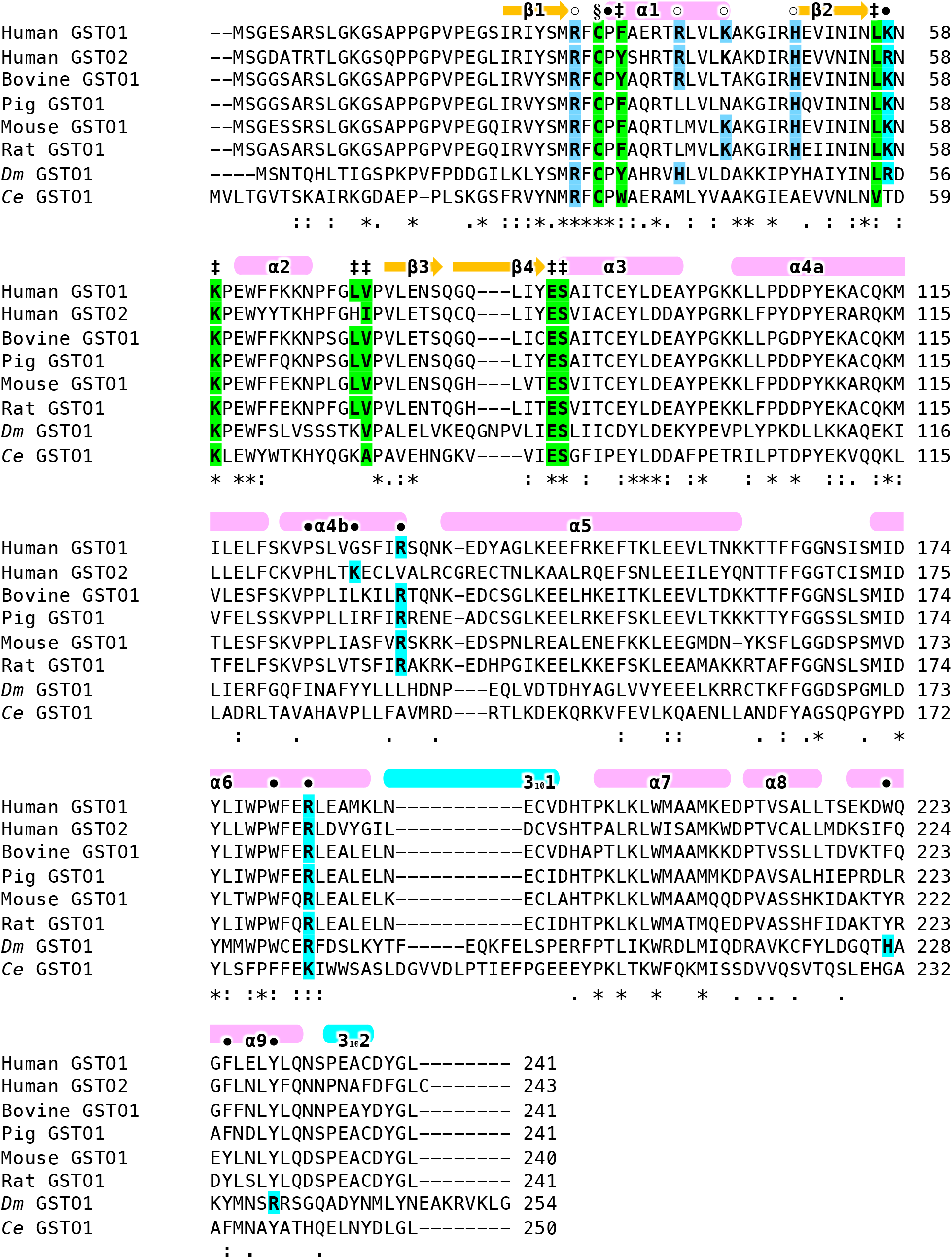
Alignment of representative GSTO1 sequences. The sequences and their UniProt IDs are as follows: Human GSTO1, P78417; human GSTO2, Q9H4Y5; Bovine GSTO1, F1MKB7; Pig GSTO1, Q9N1F5; Mouse GSTO1, O09131; Rat GSTO1, Q9Z339; *Dm* GSTO1, Q9VSL6 (*Drosophila melanogaster*); *Ce* GSTO1, P34345 (*Caenorhabditis elegans*). The secondary structure of human GSTO1 is indicated above the sequences. The catalytic cysteine is indicated by a section sign (§). G-site residues are indicated by a double daggers (‡) and are highlighted in green when identical to the human GSTO1 sequence. H-site residues are indicated by a closed circle (•) and are highlight in cyan when basic. Accessory site residues are indicated by open circles (°) and are highlight in sky blue when basic.

All GSTO homologs contain cysteine residues equivalent to C32 in GSTO1 and O2. A common feature of the H-sites is the presence of basic residues. These appear to contribute to the positively charged surfaces in the H-sites of these models (Supplementary Figure 2). All GSTO homologs considered here have a residue equivalent to R183 in human GSTO1 (Figure 2, 3). Residue R132 is not conserved in GSTO2 or in the *Drosophila* or *Caenorhabditis* homologs.

## Discussion

The calculations presented here support MMA^V^ binding in the H-site of hGSTO1-1 for reduction. This is consistent with reductase activity being competitively inhibited by 1-chloro-2,4-dinitrobenzene (CDNB) (Zakharyan et al., 2001), a ligand known to bind in the H-site of several classes of GST (although not observed in the active site of GSTO1 to date). The data suggest that two local minima exist for MMA^V^ binding, and that both involve the formation of a salt bridge with the guanidinium group of R132 (Figure 1C, D).

While the MD data indicate that MMA^V^ is likely to be loosely bound at R132, it should be noted that structural stability is not necessary for thermodynamic stability or catalysis. This contention is supported by the observation that R132 is not conserved in hGSTO2-2 which has similar MMA^V^ reductase activity to hGSTO1-1 (Schmuck et al., 2005). Arsenic’s affinity for sulfur is long-known. Mono- and dimethylarsonate species react with cysteine and GSH in aqueous solution to produce species with two or three thiol ligands (Cullen et al., 1984). They proposed the following reactions:

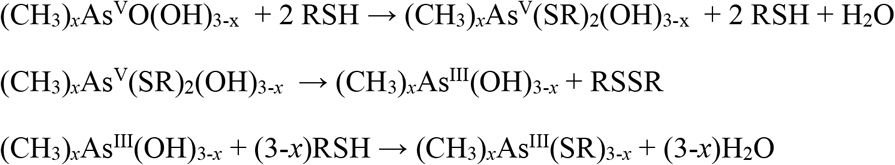

(*x* = 1, 2; RSH = cysteine or GSH)

Taken together, these observations lead us to propose a mechanism (Figure 4) for GSTO1-1 catalysed MMA^V^ reduction. Initially, GSH and MMA^V^ bind in the G- and H-sites respectively (①). Next, nucleophilic attack of GSH on MMA^V^ leads to formation of intermediate CH_3_As^V^(SG)OOH (②). The formation of the intermediate CH_3_As^V^(SG)OOH in the active site of GSTO1 is not expected to lead to the formation of the disulfide GSTO1-C32Sγ- (CH_3_)As^V^(SG)OH since C32-SγH (buried under GSH) would be sterically inaccessible.

**Figure 4.**
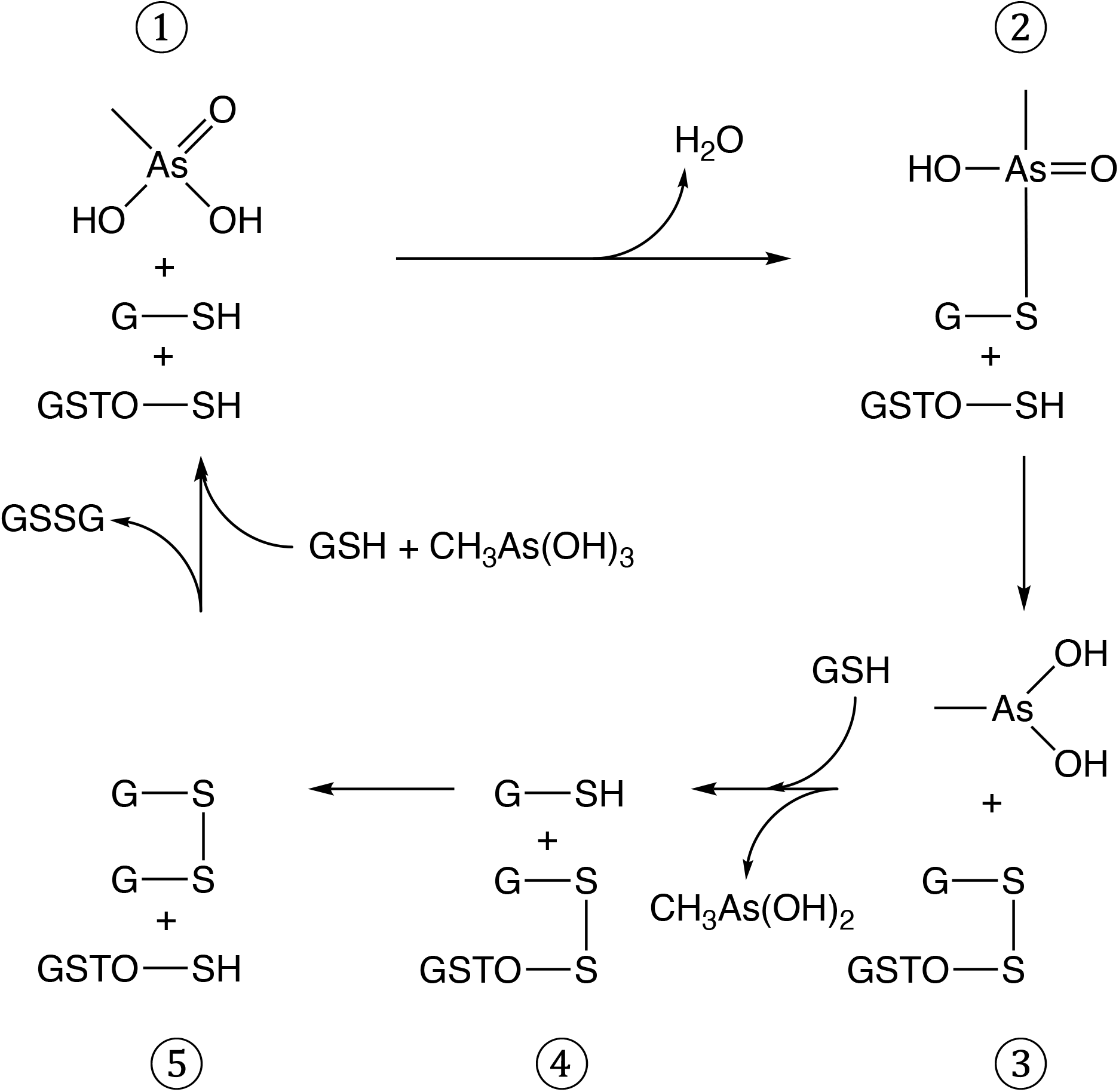
Proposed reaction mechanism for MMA^V^ reductase activity in omega-class GSTs. Intermediates ① to ⑤ in the cycle are described in the text.

Instead, C32-SγH can engage in nucleophilic attack on the GSH conjugate to release CH_3_As^III^O(OH) and give the oxidised enzyme GSTO1-S-SG (③). The latter can be reduced by a second GSH molecule (④) to give GSTO1 and GSSG (⑤). GSTO1-SSG and GSTO1 bound to GSSG have been observed in crystal structures (Board et al., 2000, Brock et al., 2013). The net reaction is thus CH_3_As^V^O(OH)_2_ + 2GSH → CH_3_As^III^(OH)_2_ + H_2_O + GSSG.

The reaction mechanism proposed here exhibits differences with respect to that proposed for the ArsC arsenate reductase, which has been studied extensively (Rosen et al., 2020). This enzyme reduces As^V^O(OH)_3_ to As^III^(OH)_3_ with the concomitant oxidation of glutaredoxin (Grx) and GSH to Grx-S-SG. Intriguingly, ArsC (like the GSTs) contains a thioredoxin domain and utilizes a catalytic cysteine residue (C12) in a position structurally equivalent to C32 in GSTO1. However, the region in ArsC equivalent to the G-site in GSTO1 would not appear to allow GSH-binding as GSH would sterically clash with the sidechains of R60, R94 and R107. ArsC contains a unique α-helical domain inserted between strands β2 and β2 of the thioredoxin domain that contains R60, which assists in substrate recognition (DeMel et al., 2004). Unlike the mechanism proposed here for GSTO1, the proposed ArsC reduction cycle includes the arsenic disulfide ArsC-S-As^V^(SG)O(OH).

Comparison of the crystal structures of hGSTO1, hGSTO2 and recently released AlphaFold models of GSTO homologs from mouse, rat, *D. melanogaster* and *C. elegans* (Figure 2A–F) allow comparison of features between widely diverged members of the omega class. All key secondary-structure elements are conserved. The AlphaFold models of mouse, rat, *D. melanogaster* and *C. elegans* GSTO appear to have G-sites compatible with GSH binding in the manner observed in the crystal structures of hGSTO1-1 and O2-2 and the presence of cysteine residues equivalent to C32 in the human GSTOs strongly implies catalysis of similar GSH-dependent reduction reactions. While H-sites show more diversity, all appear to be predominantly basic, suggesting that they bind negatively charged co-substrates. The *C. elegans* model has the smallest and least basic H-site (Figure 2F, Supplementary Figure 2F). While the MD data suggests a key role for R132 in binding MMA^V^, comparison of the H-sites reveals that while R132 is conserved in the mouse and rat homologs, it is not in hGSTO2, *D. melanogaster* or *C. elegans* homologs. As hGSTO2 also catalyses MMAV reduction, The function of R132 could be substituted in hGSTO2 by K128, located 1 turn away on helix α4b relative to R132 in GSTO1. In *Drosophila*, the function of R132 could be replaced by R234 in helix α9. Intriguingly, all models contain a tyrosine residue on helix α9 in the H-site: Y229/Y230 in hGSTO1 and hGSTO2, Y228 in mouse, Y225 in rat, Y230 in *D. melanogaster* and Y238 in *C. elegans*. Tyrosine residues in the H-sites of some GST isozymes have been shown to participate directly in catalysis. In human GSTP1-1, Y108 participates in Michael addition reactions by donating a Hydrogen-bond to co-substrates (LoBello et al., 1997). The rat GSTM2-2 H-site residue Y115 was proposed to act as an electrophile to stabilize the transition state for the epoxide ring-opening reaction and the stereospecific formation of (9*S*,10*S*)-9-(S-glutathiony1)-10-hydroxy-9,10-dihydrophenanthrene (Ji et al., 1994). In the simulations, the Y229-Oη group occasionally formed a hydrogen bond with the GSH sulfhydryl group and may play a role in activating this substrate in a manner analogous to activation in GSTs containing catalytic serine or tyrosine residues. It may potentially assist catalysis by forming a hydrogen bond with MMA^V^ and its GSH-conjugate.

It is noteworthy that arsenic-based compounds continue to attract interest as therapeutics (Chen et al., 2015). A remarkable example is the curing of acute promyelocytic leukemia (APL) patients by the combination of retinoic acid (RA) and arsenic trioxide (de Thé et al., 2017). Individual differences in response to arsenic, including fatal adverse reactions during ATO therapy, have been reported. Further understanding of arsenic metabolizing enzymes such as GSTO1 and GSTO2 will assist in the development of arsenic-based therapeutics.

## Acknowledgements

This work was supported by Project Grants APP1124673 and APP1156455 from the National Health and Medical Research Council of Australia (NHMRC). Molecular graphics and analyses performed with UCSF ChimeraX, developed by the Resource for Biocomputing, Visualization, and Informatics at the University of California, San Francisco, with support from National Institutes of Health R01-GM129325 and the Office of Cyber Infrastructure and Computational Biology, National Institute of Allergy and Infectious Diseases. The author would like to thank Prof. Philip Board for helpful discussions.

---

**Supplementary Figure 1.**
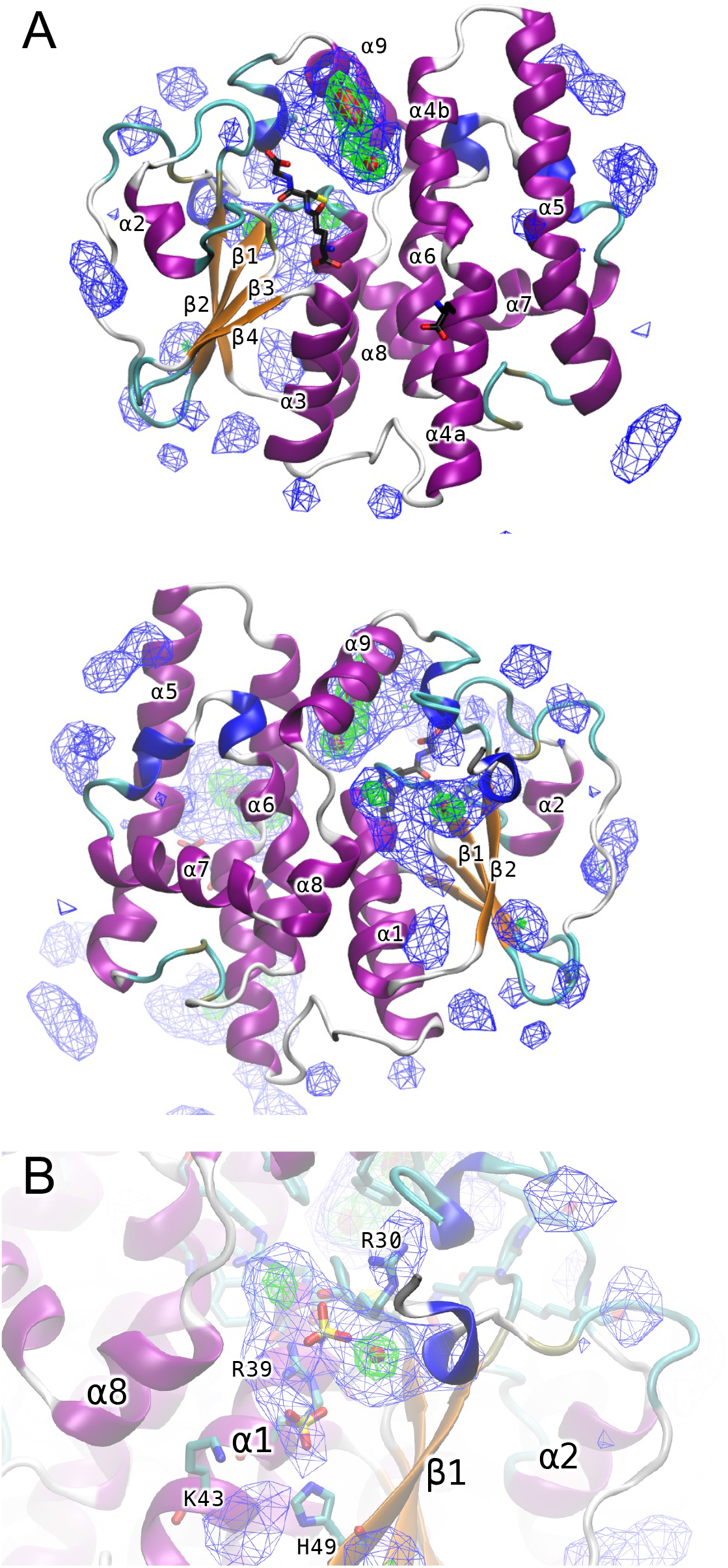
(A) Two views of the GSTO1 monomer in cartoon form rotated by 180° vertically. (B) Accessory binding site. The location of sulfate ions from PDB 5YVN are shown for comparison. Density isosurfaces for P atoms in blue, green and red are contoured at 0.02, 0.1 and 0.2 Da Å^−3^ respectively.

**Supplementary Figure 2.**
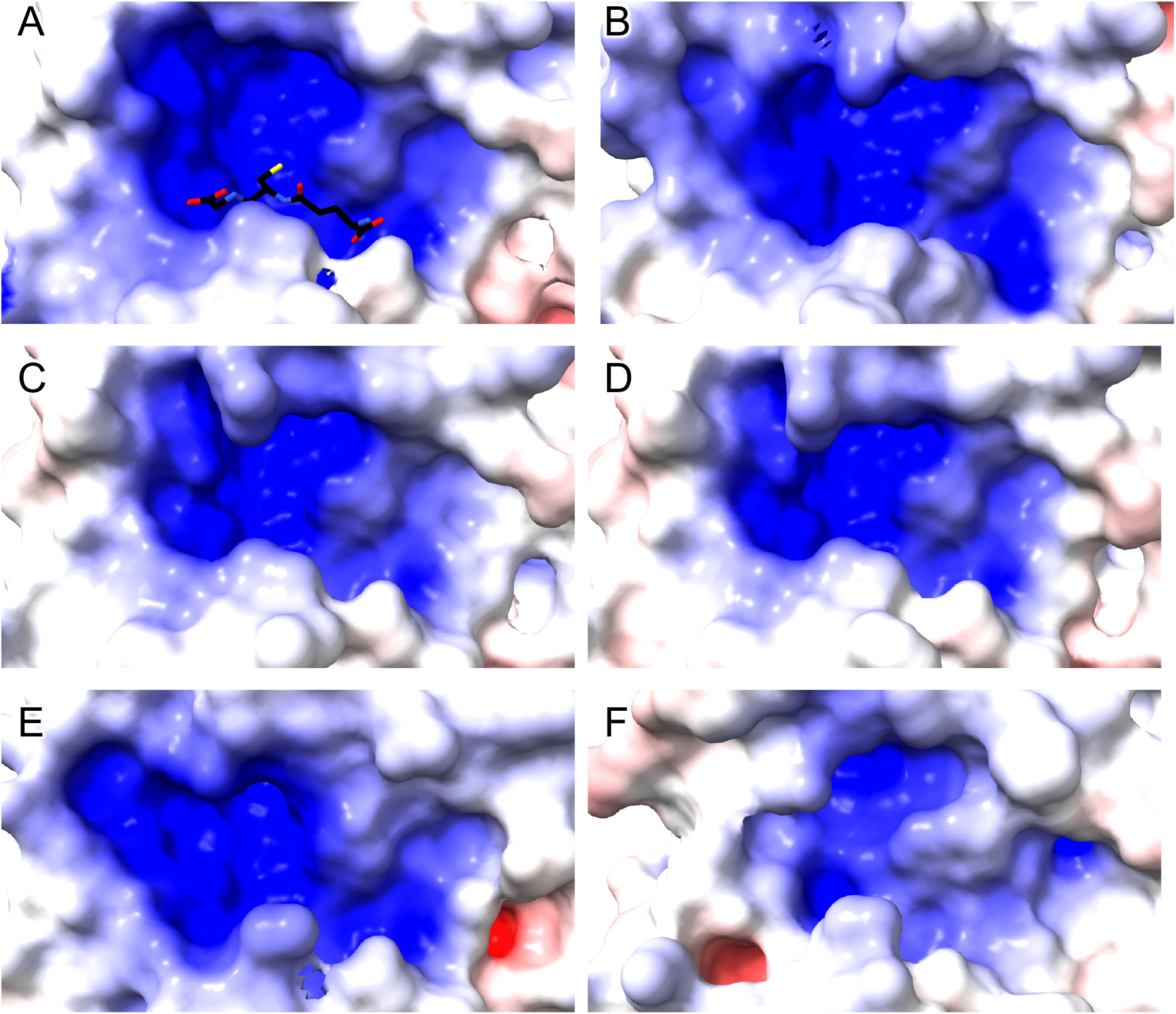
Surface representations of GSTO homologs coloured by electrostatic potential ranging from −8 kT/e (red) to +8 kT/e (blue). (A) human GSTO1 (PDB ID 1EEM) with GSH in stick form. (B) human GSTO2 (PDB ID 3Q19) and AlphaFold models of (C) mouse GSTO1, (D) rat GSTO1, (E) *Drosophila melanogaster* GSTO1, and (F) *Caenorhabditis elegans* GSTO1.

