## Supplementary Figures for "Proposed Mechanism for Monomethylarsonate Reductase Activity of Human Omega-class Glutathione Transferase GSTO1-1"

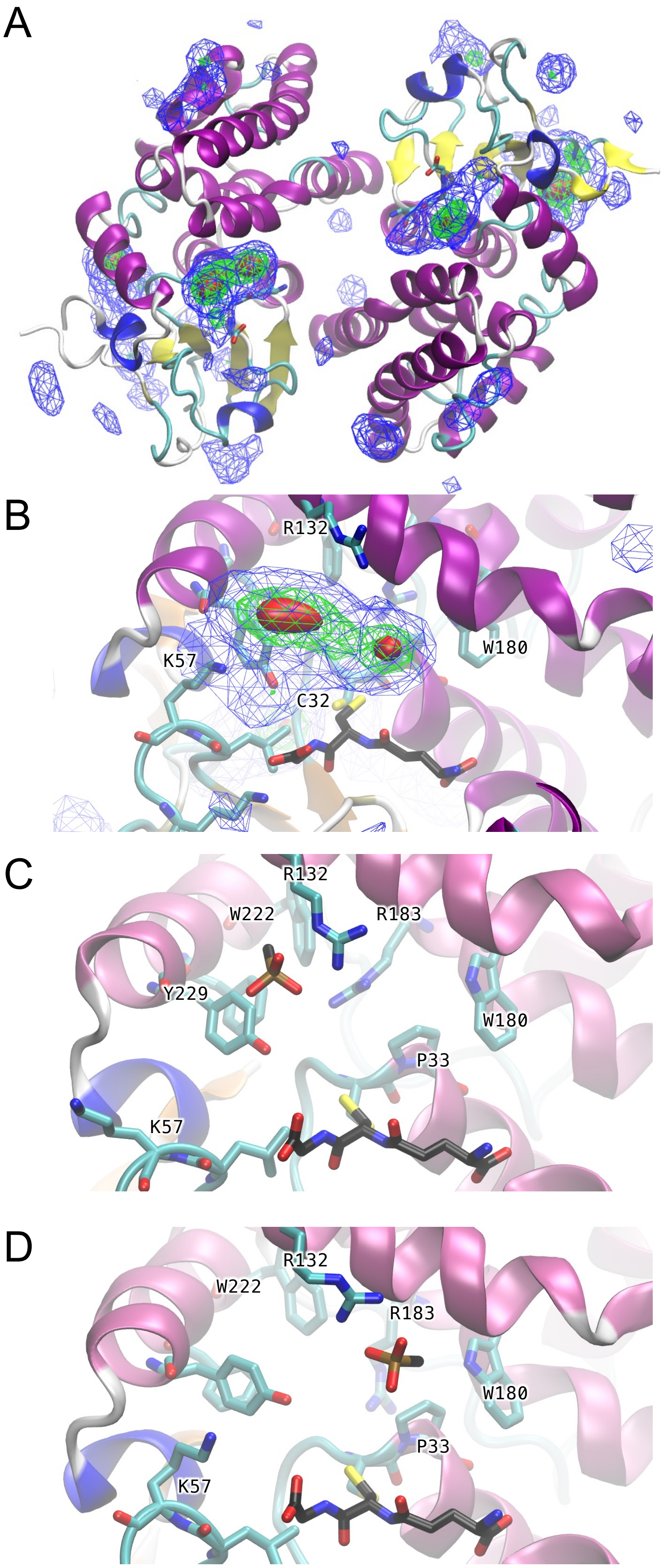
A
B
R132
W180
K57
C32
C
R132
W222
R183
Y229
W180
P33
K57
D
R132
W222
R183
W180
P33
K57

Figure 1

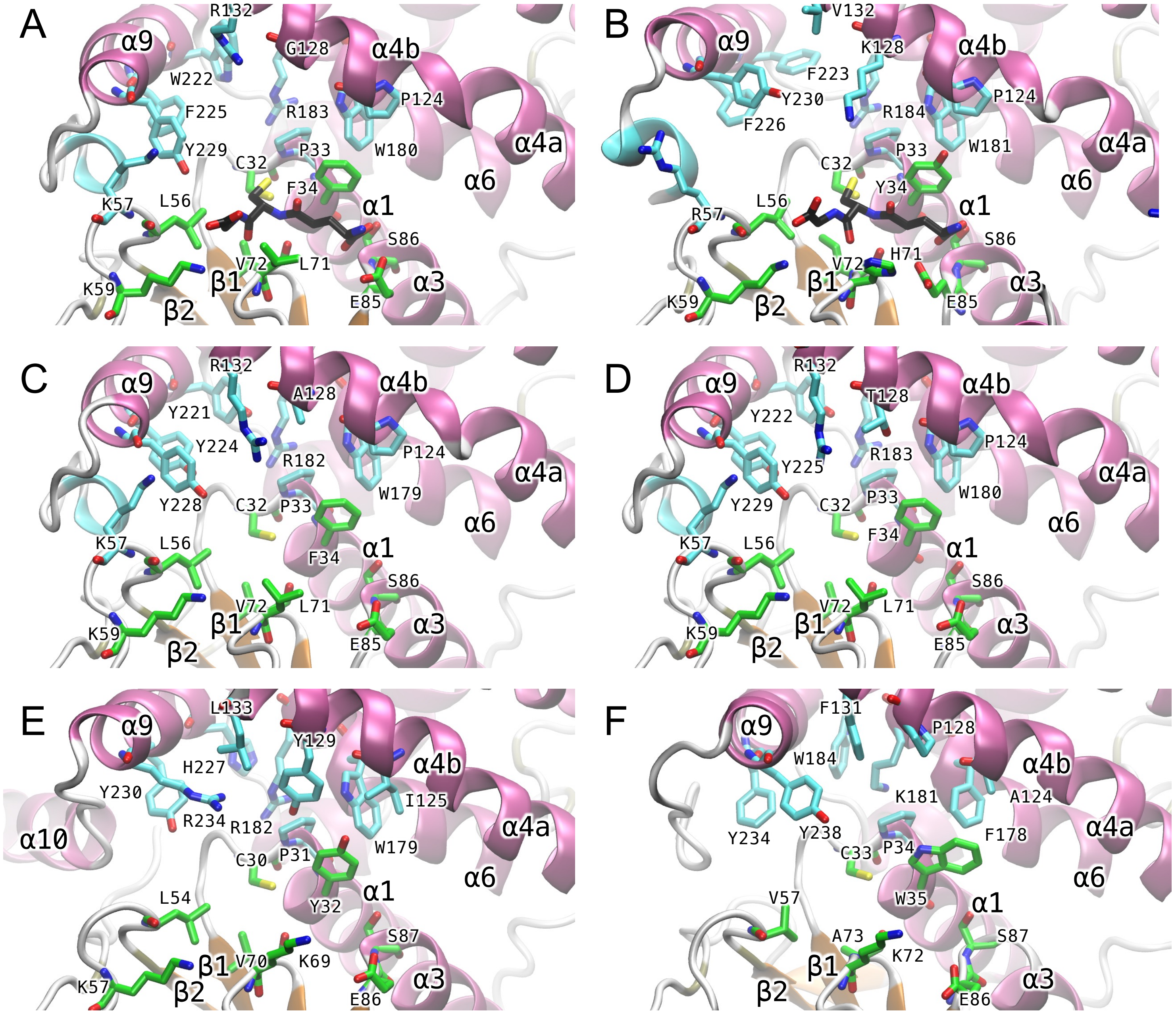
A
α9
R132
G128
α4b
W222
F225
R183
P124
Y229
C32
P33
W180
α4a
α6
K57
L56
F34
α1
S86
K59
V72
L71
β1
α3
β2
E85
B
V132
α9
K128
α4b
F223
Y230
R184
P124
F226
W181
α4a
C32
P33
α6
Y34
R57
L56
α1
S86
H71
V72
K59
β1
α3
β2
E85
C
α9
R132
A128
α4b
Y221
Y224
R182
P124
α4a
W179
Y228
C32
P33
α6
F34
K57
L56
α1
F34
S86
V72
L71
β1
α3
K59
β2
E85
D
α9
R132
T128
α4b
Y222
P124
R183
Y225
α4a
W180
Y229
C32
P33
α6
F34
K57
L56
α1
S86
V72
L71
β1
α3
K59
β2
E85
E
α9
L133
Y129
α4b
H227
I125
Y230
R234
R182
α4a
α10
W179
C30
P31
α6
Y32
α1
L54
S87
β1
V70
K69
α3
K57
β2
E86
F
α9
F131
P128
W184
α4b
K181
A124
Y234
Y238
F178
α4a
C33
P34
α6
W35
α1
V57
A73
S87
β1
K72
α3
β2
E86

Figure 2

```
                                        β1  ○ §•‡ α1 ○   ○     ○ β2   ‡•
Human GST01  --MSGESARSLGKGSAPPGPVPEGSIRIYSMRFCPFAERTRLVLKAKGIRHEVININLKN  58
Human GST02  --MSGDATRTLGKGSQPPGPVPEGLIRIYSMRFCPYSHRTRLVLKAKDIRHEVVNINLRN  58
Bovine GST01 --MSGGSARSLGKGSAPPGPVPEGLIRVYSMRFCPYAQRTRLVLTAKGIRHEVININLKN  58
Pig GST01    --MSGGSARSLGKGSAPPGPVPEGLIRVYSMRFCPFAQRTLLVLNAKGIRHQVININLKN  58
Mouse GST01  --MSGESSRSLGKGSAPPGPVPEGQIRVYSMRFCPFAQRTLMVLKAKGIRHEVININLKN  58
Rat GST01    --MSGASARSLGKGSAPPGPVPEGQIRVYSMRFCPFAQRTLMVLKAKGIRHEIININLKN  58
Dm GST01     ----MSNTQHLTIGSPKPVFPDDGILKLYSMRFCPYAHRVHLVLDAKKIPYHAIYINLRD  56
Ce GST01     MVLTGVTSKAIRKGDAEP-PLSKGSFRVYNMRFCPWAERAMLYVAAKGIEAEVVNLNVTD  59
                    :: :  *.  *    .* :::*.*****::.*. : : ** *  . : :*: :

             ‡    α2     ‡‡   β3       β4 ‡‡   α3              α4a
Human GST01  KPEWFFKKNPFGLVPVLENSQGQ---LIYESAITCEYLDEAYPGKKLLPDDPYEKACQKM 115
Human GST02  KPEWYYTKHPFGHIPVLETSQCQ---LIYESVIACEYLDDAYPGRKLFPYDPYERARQKM 115
Bovine GST01 KPEWFFKKNPSGLVPVLETSQGQ---LICESAITCEYLDEAYPGKKLLPGDPYEKACQKM 115
Pig GST01    KPEWFFQKNPSGLVPVLENSQGQ---LIYESAITCEYLDEAYPGKKLLPDDPYEKACQKM 115
Mouse GST01  KPEWFFEKNPLGLVPVLENSQGH---LVTESVITCEYLDEAYPEKKLFPDDPYKKARQKM 115
Rat GST01    KPEWFFEKNPFGLVPVLENTQGH---LITESVITCEYLDEAYPEKKLFPDDPYEKACQKM 115
Dm GST01     KPEWFSLVSSSTKVPALELVKEQGNPVLIESLIICDYLDEKYPEVPLYPKDLLKKAQEKI 116
Ce GST01     KLEWYWTKHYQGKAPAVEHNGKV----VIESGFIPEYLDDAFPETRILPTDPYEKVQQKL 115
             * **:         *.:*         : ** :  :***: :*   : * *  ::. :*:

                     •α4b•   •              α5
Human GST01  ILELFSKVPSLVGSFIRSQNK-EDYAGLKEEFRKEFTKLEEVLTNKKTTFFGGNSISMID 174
Human GST02  LLELFCKVPHLTKECLVALRCGRECTNLKAALRQEFSNLEEILEYQNTTFFGGTCISMID 175
Bovine GST01 VLESFSKVPPLILKILRTQNK-EDCSGLKEELHKEITKLEEVLTDKKTTFFGGNSLSMID 174
Pig GST01    VFELSSKVPPLLIRFIRRENE-ADCSGLKEELRKEFSKLEEVLTKKKTTYFGGSSLSMID 174
Mouse GST01  TLESFSKVPPLIASFVRSKRK-EDSPNLREALENEFKKLEEGMDN-YKSFLGGDSPSMVD 173
Rat GST01    TFELFSKVPSLVTSFIRAKRK-EDHPGIKEELKKEFSKLEEAMAKKRTAFFGGNSLSMID 174
Dm GST01     LIERFGQFINAFYYLLLHDNP---EQLVDTDHYAGLVVYEEELKRRCTKFFGGDSPGMLD 173
Ce GST01     LADRLTAVAHAVPLLFAVMRD---RTLKDEKQRKVFEVLKQAENLLANDFYAGSQPGYPD 172
               :    .       .   .               :   ::      . : .*   .  *

             α6   •  •                  3_10 1      α7        α8        •
Human GST01  YLIWPWFERLEAMKLN-----------ECVDHTPKLKLWMAAMKEDPTVSALLTSEKDWQ 223
Human GST02  YLLWPWFERLDVYGIL-----------DCVSHTPALRLWISAMKWDPTVCALLMDKSIFQ 224
Bovine GST01 YLIWPWFERLEALELN-----------ECVDHAPTLKLWMAAMKKDPTVSSLLTDVKTFQ 223
Pig GST01    YLIWPWFERLEALELN-----------ECIDHTPKLKLWMAAMMKDPAVSALHIEPRDLR 223
Mouse GST01  YLTWPWFQRLEALELK-----------ECLAHTPKLKLWMAAMQQDPVASSHKIDAKTYR 222
Rat GST01    YLIWPWFQRLEALELN-----------ECIDHTPKLKLWMATMQEDPVASSHFIDAKTYR 223
Dm GST01     YMMWPWCERFDSLKYTF-----EQKFELSPERFPTLIKWRDLMIQDRAVKCFYLDGQTHA 228
Ce GST01     YLSFPFFEKIWWSASLDGVVDLPTIEFPGEEEYPKLTKWFQKMISSDVVQSVTQSLEHGA 232
             *: :*: :::                     . * *  *   *  . .. .   .

              • α9•      3_10 2
Human GST01  GFLELYLQNSPEACDYGL-------- 241
Human GST02  GFLNLYFQNNPNAFDFGLC------- 243
Bovine GST01 GFFNLYLQNNPEAYDYGL-------- 241
Pig GST01    AFNDLYLQNSPEACDYGL-------- 241
Mouse GST01  EYLNLYLQDSPEACDYGL-------- 240
Rat GST01    DYLSLYLQDSPEACDYGL-------- 241
Dm GST01     KYMNSRRSGQADYNMLYNEAKRVKLG 254
Ce GST01     AFMNAYATHQELNYDLGL-------- 250
              : .     .
```

Figure 3

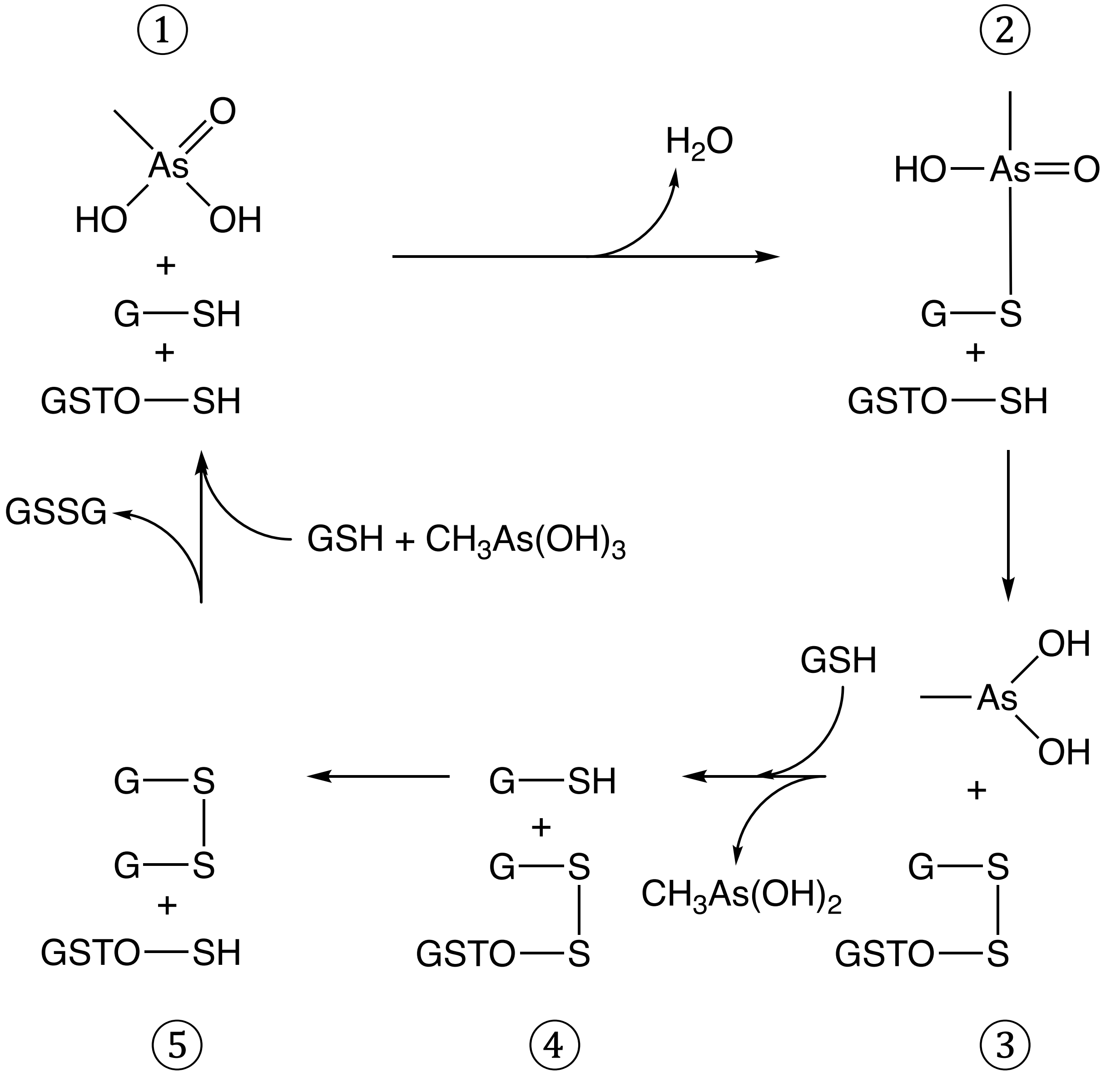
1
2
3
4
5
H2O
HO—As=O
G—S
GSTO—SH
G—SH
GSSG
GSH + CH3As(OH)3
GSH
OH
As
CH3As(OH)2
GSTO—S

Figure 4

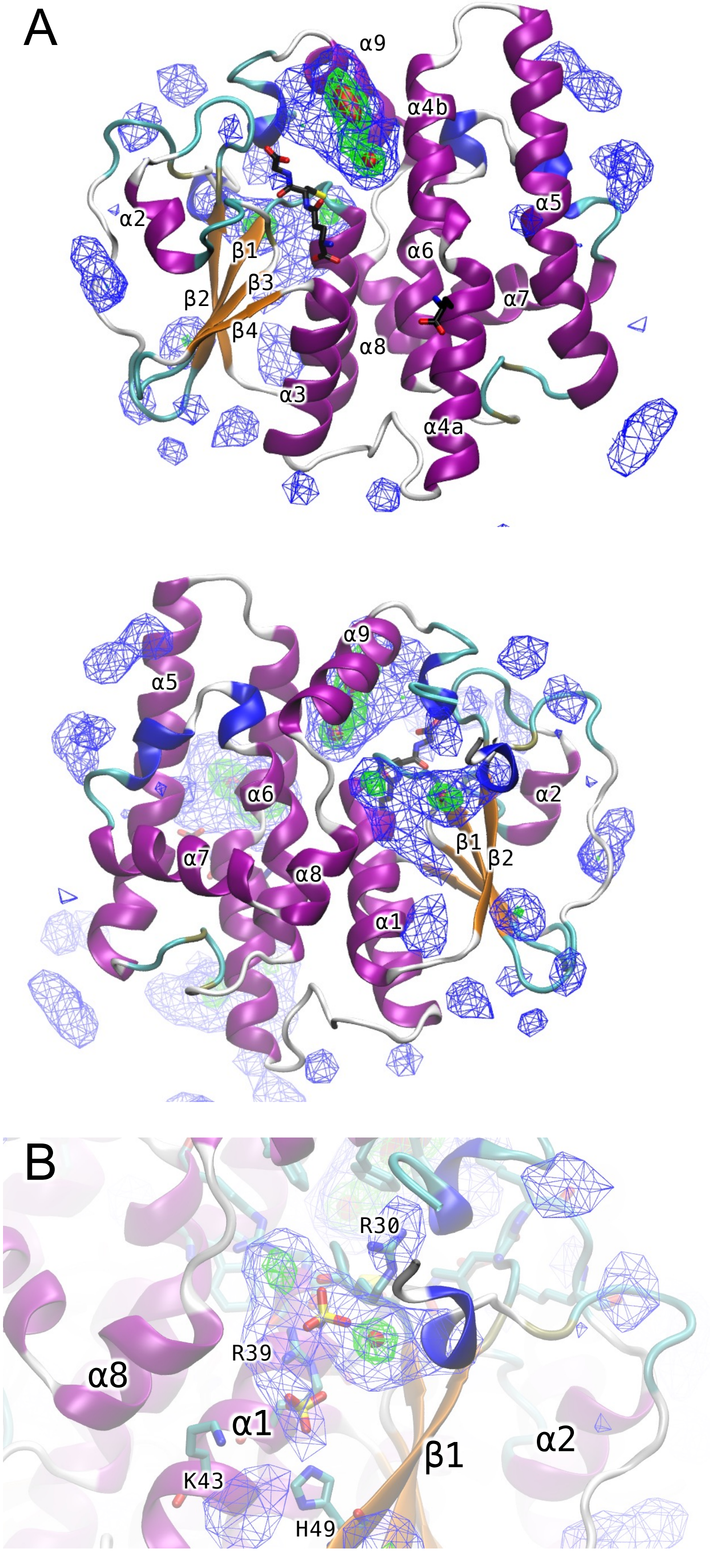


Supplementary Figure 1

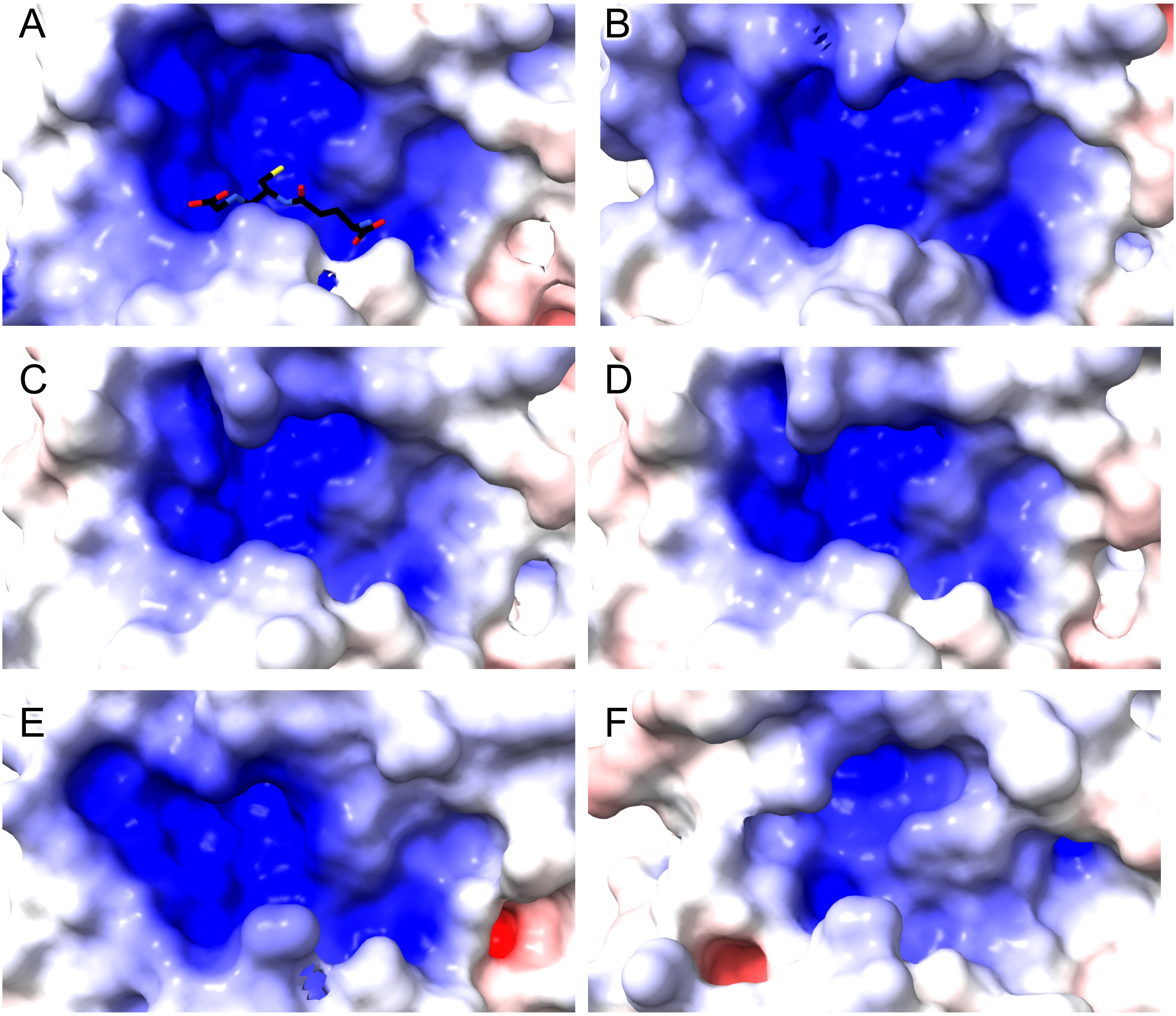


Supplementary Figure 2
